# Accelerated amyloid neurodegeneration in HIV-1-infected APP-KI Alzheimer’s disease mice

**DOI:** 10.64898/2026.05.16.725620

**Authors:** Shaurav Bhattarai, Emma G. Foster, Rana Kadry, Yaman Lu, Mohit Kumar, Syed Qasim, Amrita Mitra, Harsh Pathak, Larisa Y. Poluektova, Santhi Gorantla, R. Lee Mosley, Pravin Yeapuri, Howard E. Gendelman

## Abstract

**INTRODUCTION:** A higher incidence of dementia, including Alzheimer ’s-like pathology, is observed in aged people living with HIV-1. However, mechanisms linking HIV-1 to Alzheimer’s disease (AD) pathology remain unclear, due to the lack of animal models that allow for concurrent study.

**METHODS:** We created a novel APP knock-in (KI) AD mouse, NOG/APP^KM670,671NL^/IL-34 (hNAIL) that permits study of progressive brain HIV-1 replication. The mice harbor human microglia-like cells. Four-month-old CD34+ human cell reconstituted mice infected with the HIV-1_ADA_ strain facilitated studies of HIV-1 replication on AD pathologies.

**RESULTS:** HIV-1 replication increased Aβ levels and reduced synaptic and neuronal integrity. Spatial transcriptomics demonstrated distinct Aβ and HIV-1 transcriptional patterns, whereas dual diseased combinations amplified AD pathology. Neurons showed highest transcriptional change, with genes linked to neuroinflammation, protein trafficking, and synaptic dysfunction.

**DISCUSSION:** The hNAIL mice enable interrogation of HIV-AD comorbidities, with a future potential for the development of novel therapeutic interventions.

## 1. Background

Widespread access to antiretroviral therapy (ART) has transformed human immunodeficiency virus-1 (HIV-1) infection into a chronic condition, resulting in a growing aging population of people living with HIV-1 (PLWH) [1-3]. Despite effective viral suppression by ART, cognitive impairments remain prevalent in PLWH [4], and diagnostic frameworks such as the Frascati criteria continue to evolve to improve classification and enable cross-cohort comparisons [5]. Emerging clinical and preclinical studies suggest that HIV-1 infection is associated with an increased risk of AD and related dementias [6-8]. The accumulation of AD-associated proteins, including amyloid-β (Aβ) and phosphorylated tau (p-tau), is frequently observed in aged PLWH [9-11].

Neuroinflammation drives both HIV-associated neurocognitive disorders (HAND) and AD [12, 13]. Microglia regulate inflammation and serve as a major reservoir for persistent central nervous system (CNS) HIV-1 infection [14]. In parallel, microglial responses to amyloid plaques and tau pathology critically shape the progression of AD [15]. Thus, microglia represent a central intersection between viral persistence and neurodegeneration. Mechanistic studies have further demonstrated that HIV-1 proteins, such as Tat, interact with Aβ and tau, impairing Aβ clearance and exacerbating neurodegenerative processes [16, 17].

Despite the convergent pathology, the mechanistic understanding of HIV-AD interactions remains limited. Existing preclinical AD models are developed on immunocompetent backgrounds and cannot support sustained HIV-1 infection, as the virus is species-restricted and requires human immune cells for productive replication [18-20]. Conversely, current HIV-1 models lack AD-relevant pathology because mice do not naturally develop AD. Commonly used AD transgenic mouse models rely on overexpression of familial AD mutations (FAD) under heterologous promoters to accelerate pathology [21, 22]. While these models have been instrumental in dissecting aspects of AD pathology, they do not recapitulate the slow, progressive nature of AD-like diseases in PLWH. Importantly, in ART-treated PLWH, one of the key clinical questions is whether chronic HIV-1-associated inflammation and comorbidities modulate early AD-like changes rather than end-stage Alzheimer’s dementia [23]. Additionally, autopsy studies in PLWH frequently report low-to-intermediate AD neuropathological changes, with predominantly diffuse plaques and intraneuronal amyloid burden, highlighting the need for models that capture gradual, preclinical-stage pathology [11, 24].

In recent years, knock-in (KI) AD models have addressed some of these limitations by incorporating familial mutations into the endogenous mouse *App* locus, thereby avoiding overexpression artifacts [25, 26]. However, the immunocompetent background of these models precludes engraftment of human hematopoietic stem cells (HSCs), limiting HIV-1 studies [18, 20]. Alternative experimental platforms, including human brain organoids, peripheral blood leukocyte (PBL)-humanized mice, and xenograft microglial models, have been used to study HIV and AD interactions [27-31]. However, each of these platforms lacks sustained progressive HIV-1 replication or fails to express human AD pathological proteins. Organoid models lack the vasculature and systemic immune microenvironment required for long-term studies. Human PBL (hu-PBL) models can sustain HIV-1 infection only transiently before developing graft-versus-host disease (GvHD), limiting the experimental duration and interpretation of “true” disease-associated neuroinflammatory responses. Xenograft-based approaches that transplant human glial cells into the rodent brain allow HIV-1 brain infection, but lack systemic human immune interactions and a relevant microglia population [18, 20, 32].

To address these challenges, we previously developed an APP knock-in AD model (NA) with the Swedish AD mutation on an immunodeficient NOG background, which mimics early-stage AD pathology, by predominantly developing intraneuronal amyloid pathology [33]. To support brain HIV-1 replication, the NA mice were crossed with transgenic human interleukin-34 (hIL34) mice to generate the hNAIL mouse model, which supports human microglia-like cells in the brain following HSC engraftment [33-35].

HSCs reconstituted hNAIL mice were infected with macrophage-tropic HIV-1_ADA_ strain at 4 months, and sacrificed at 6 months to evaluate the role of sustained HIV-1 infection in AD pathology. We found that HIV-1 infection potentiated early AD-like pathologies, including increased amyloid accumulation, microglial activation, and synaptic injury. The hNAIL mouse model serves as a unique platform for studying HIV-AD comorbidities.

## 2. Methods

### 2.1 Study approvals

All animal experimental procedures were approved by the Institutional Animal Care and Use Committee (IACUC) of the University of Nebraska Medical Center (UNMC) (18-109-08). All animal procedures were conducted in accordance with the National Institutes of Health Guide for the Care and Use of Laboratory Animals. The University of Nebraska Medical Center is an Association for Assessment and Accreditation of Laboratory Animal Care-accredited institution.

### 2.2 Generation of hNAIL mice, humanization, and HIV-1 infection

hNAIL mice were generated as previously described [33, 34]. Briefly, a single amyloid precursor protein (APP) KI mutation (NA) was introduced carrying the Swedish familial AD mutation APP^KM670,671NL^ on an immunodeficient NOG (NOD.Cg-Prkdc^scid^Il2rg^tm1Sug^/JicTac) background. These animals were then crossed with NOG mice carrying the IL-34 transgene, NOG (NOD.Cg-*Prkdc ^scid^ Il2rg ^tm1Sug^* Tg(CMV*-IL-34*)*1*/Jic). These animals develop human microglia-like cells after hematopoietic stem cell (HSC) reconstitution [34, 35]. hNAIL mice were homozygous for the APP KI gene and heterozygous for the human *IL34* transgene.

For human immune cell engraftment, 2-3-day-old pups were irradiated with 1 Gy (RS 2000 X-ray Irradiator). Four hours after irradiation, the pups were intrahepatically injected with 50,000 CD34+ HSCs derived from cord blood (STEMCELL Technologies, Vancouver, Canada). Mice were bled at 4 months of age and analyzed by flow cytometry to confirm human cell engraftment.

Four-month-old, HSC engrafted hIL34 (n=9) and hNAIL mice (n=9) were infected intraperitoneally with 10^4^ TCID_50_ of the HIV-1_ADA_ viral strain. HSC-reconstituted hNAIL (n=9) and NOG mice (n=7) served as controls. Mice were bled 14 days post-infection to measure the plasma viral load using the automated COBAS Ampliprep V2.0/Taqman-48 system (Roche, Boston, MA) [36]. The mice were euthanized eight weeks post-infection. The brain, blood, and spleen were collected for further analysis. The right hemisphere of the brain was used for histology, and the left hemisphere was used for biochemical assays.

### 2.3 Flow cytometry

Blood samples were obtained via submandibular bleeding into EDTA-treated tubes (BD Microtainer, Franklin Lakes, NJ, USA) and centrifuged at 350 × g for 10 min. The resulting blood cells were resuspended in a FACS buffer containing 2% fetal bovine serum (FBS) and 0.1% sodium azide in phosphate-buffered saline (PBS). Cells were then incubated at 4°C for 30 min with an antibody cocktail containing human-specific FITC-anti-CD45 (BD Bioscience, cat. 555482), AF700-anti-CD3 (BD Bioscience, cat. 557943), PE-Cy5-anti-CD19 (BD Bioscience, cat. 555414), APC-anti-CD4 (BD Bioscience, cat. 555349), BV421-anti-CD8 (BD Bioscience, cat. 562428) and PE-anti-CD14 (BD Bioscience, cat. 555398). Red blood cells were lysed with FACS lysing solution (BD Biosciences, cat.349202). After lysis, the cells were washed with 2 mL FACS buffer and fixed in 2% paraformaldehyde. Data acquisition was performed using FACS Diva v6 software (BD Biosciences) on a BD LSR2 flow cytometer, and data analysis was conducted using FlowJo v10.2 (Tree Star, Ashland, OR, USA). The gating strategies were determined based on appropriate controls.

### 2.4 Immunofluorescence (IF) and Immunohistochemistry (IHC)

During necropsy, the right hemisphere of the mouse brain was fixed using depolymerized 4% paraformaldehyde for 24 h at 4°C. The fixed brains were processed overnight using an Epredia STP 120 Spin Tissue Processor (Thermo Fisher Scientific). Following processing, the tissues were embedded in paraffin blocks and stored at 4°C until sectioning. Five-micrometer-thick sagittal tissue sections were prepared and collected on pre-treated microscope slides (Fisher Scientific, cat. 1255015) for IHC and IF staining, respectively. Tissue slides were first deparaffinized and rehydrated, and antigen retrieval was performed using a Trilogy solution (Sigma-Aldrich, cat. 920P-07) by heat and pressure induction [35]. Brain sections were blocked with 10% normal goat serum to prevent nonspecific staining. The sections were incubated with a primary mouse antibody against human Aβ using the 6E10 antibody (BioLegend, San Diego, CA, cat. 803001), HIV-1p24 (Santa Cruz Biotechnology, Dallas, TX, cat.SC-65918), rabbit antibody against human TMEM-19 (Abcam, Cat.185333), or human P2RY12 (Abcam, Cat. HPA013796) overnight at 4°C. After three washes, the tissue sections were incubated with HRP-conjugated anti-mouse-rabbit-rat secondary antibody (GeneTex, Irvine, CA, cat. GTX83397) for 1 h at room temperature. For chromogenic detection, 3,3’ diaminobenzidine (DAB) (Sigma-Aldrich, cat. D4239) was used, followed by hematoxylin (VWR, Radnor, PA, Cat. 10015-074, VWR) for nuclear counterstaining purposes. Slides were imaged using a Zeiss Axioscan 7 whole-slide imaging system.

For IF staining, the slides were stained with a rabbit primary antibody against synaptophysin (Invitrogen, cat. PA1-1043) and mouse antibody against PSD-95 (Invitrogen, cat. MA1-046) or rabbit antibody against human IBA1 (Abcam, cat. AB221933-1001) and HIV-1p24 (Santa Cruz Biotechnology, cat.SC-65918) or mouse primary antibody against MAP2 (Invitrogen, cat. 13-1500) and guinea pig antibody against Neun (Synaptic system, cat. 266004) or rabbit anti IBA1 (Fujifilm, cat.01327691) and chicken antibody against GFAP (Invitrogen, cat.PA1-10004) and incubated overnight at 4°C in a humidified chamber. Following washes, sections were incubated with Alexa 488-conjugated goat anti-rabbit IgG (Invitrogen, cat.A11008), Alexa568-conjugated goat anti-mouse IgG (Invitrogen, cat. A-11004), and Alexa594-conjugated goat anti-chicken IgG (Invitrogen, cat. A11042), and Alexa488-conjugated goat anti-mouse IgG (Invitrogen, cat. A32723) or Alexa 568-conjugated goat anti-guinea pig IgG (Invitrogen, catalog number. A11075) for one hour at room temperature. Following washing, DAPI was used for nuclear staining (Thermo Fisher Scientific, cat. D1306). Slides were imaged using the Zeiss Axioscan 7 whole slide imaging system. Images were quantified using the HALO software (v3 version) [33]. Aβ IHC was quantified using the Indica-Labs Area Quantification v2.4.3 module. For IF image analysis, the Indica-Labs Area Quantification FL v2.3.4 module was used. The percentage of stained area was used for analysis. The Indica Labs-Microglial Activation FL v1.0.6 module was used to quantify activated microglia. Microglial cells with less than 15 processes and a total process area greater than 15 µm^2^ were scored as activated microglia.

### 2.5 ELISA

Snap-frozen mouse hippocampal and cortical tissues were homogenized in ice-cold 50mM Tris-HCl buffer (pH 7.6) containing 150 mM NaCl and a protease-phosphatase inhibitor cocktail (Thermo Fisher, cat. 78445). Homogenates were centrifuged at 20,000x g for 60 minutes at 4°C, and the resulting supernatants were collected for soluble Aβ_42_ quantification. To quantify insoluble Aβ_42_, the remaining pellets were resuspended in 6M guanidine-HCl and centrifuged at 20,000 × g for 60 min at room temperature. Aβ42 levels in the brain cortex and hippocampus homogenates were quantified using an ELISA kit (R&D Systems, Waltham, MA, cat. DAB142).

### 2.6 Western blot

Frozen brain cortex tissues were processed to evaluate the protein levels of full-length APP and APP fragments CTF-ɑ and CTF-β, as described previously [37]. Samples were homogenized in 50mM Tris-HCl buffer (pH 8.0) containing 150mM NaCl, 50mM EDTA, 1% Triton X-100, and protease and phosphatase inhibitor cocktail (Thermo Fisher Scientific, cat. 78442). The homogenates were centrifuged at 20,000 × g for 1 hour at 4^0^C, and the resulting supernatants were collected. The Pierce BCA protein assay kit (Thermo Fisher Scientific, cat. 23227) was used to quantify protein concentration. Ten to fifteen micrograms of proteins were loaded onto 10% or 15% SDS-polyacrylamide gels and transferred onto activated 0.2 µm PVDF membranes. After transfer, membranes were blocked in 5% non-fat dry milk for 1 h at room temperature and then incubated with primary antibodies overnight at 4°C on a shaker. The following primary antibodies were used: anti-Aβ 1-16 antibody (1:1000, 6E10, cat. 803001), anti-APP (1:1000, clone 22C11, EMD Millipore, cat. MAB348), anti-APP C-terminal (1:1000, Millipore Sigma-Aldrich, cat.A8717), and anti-GAPDH (Cell Signaling Technology, 14C10, cat.2118S).

Following primary antibody incubation, membranes were washed three times for 10 min each with TBST containing 0.1% Tween-20. Membranes were exposed to HRP-conjugated secondary antibodies-goat anti rabbit IgG H&L (1:10,000, Abcam, cat. Ab205718) or goat anti-mouse IgG H&L (1:10,000, Abcam, cat. Ab205719) for 2 h at room temperature. Membranes were washed three times with TBST (10 minutes each) and the protein bands were visualized using either SuperSignal West Pico PLUS or SuperSignal West Femto Maximum Sensitivity Substrate (Thermo Fisher Scientific). Chemiluminescent signals were captured using an iBright 750 Imaging System, and band intensities were quantified using ImageJ software.

### 2.7 Magnetic resonance imaging (MRI)

Magnetic resonance imaging was performed on 16 mice (3 NOG, 4 HIV-1 infected hIL34, 4 hNAIL, and 5 HIV-1 infected hNAIL mice) at 8 weeks of infection (6 months of age). Imaging was performed using a Bruker 7 T scanner (PharmaScan). Mice were anesthetized with isoflurane delivered in oxygen at a flow rate of 1 L/min, and their respiration was continuously monitored throughout the procedure. High-resolution T2-weighted images were acquired using TurboRARE with repetition time/echo time (TR/TE) = 300/36 ms, RARE factor=4, number of slices =25, slice thickness=0.5 mm, field of view (FOV)= 20 × 20 mm^2,^ and matrix = 256 × 256. The cortical area was quantified using region of interest (ROI) analysis in ImageJ.

### 2.8 Behavioral tests

Memory function was assessed using novel object recognition, Y-maze, and open field tests. Behavioral data were collected at 6 months of age, approximately 8 weeks post-infection, from the NOG, HIV-1-infected hIL34 (hIL34+HIV-1), hNAIL, and HIV-1-infected hNAIL (hNAIL+HIV-1) groups. Visual recognition memory was assessed using the novel-object recognition test. The mice were habituated to a square open-field area without objects for the first 10 min. After a 90-minute rest period, the animals completed three 5-minute familiarization sessions, each separated by a 90-minute interval, during which two identical objects were presented. Ten minutes after the familiarization trial, the mice were introduced to one familiar and one novel object for a 5-minute test session. Exploratory behavior was tracked using ANY-maze software (Wood Dale, IL, USA), and the time spent investigating each object was recorded. Recognition memory was expressed as a percentage of time spent with a novel object and calculated as (time spent with novel object/total exploration time) × 100 for each animal.

The Y-maze test was performed to evaluate spatial working memory using a spontaneous alternation Y-maze paradigm, as described previously [33]. The apparatus consisted of three arms (39.5 × 8.5 × 13 cm) arranged at an angle of 120^0^ and labeled A, B, and C. Arm entries were defined as the entry of all four paws into an arm and were automatically recorded using the ANY-maze software. Spontaneous alterations were identified as sequential entries into all three arms (ABC, BAC, or CAB, but not ABA). Each mouse underwent two 8-minute sessions conducted on consecutive days, with the first serving as an acclimation trial and the second as a test trial. Data from the test trial were used to calculate the percentage of spontaneous alternations using the formula [number of spontaneous alternations/ (total number of entries-2)] x 100 for each animal.

Locomotor activity and anxiety-like behaviors were evaluated using the open-field test. Following acclimation, the mice were placed in the center of an open-field arena and allowed to explore freely for 10 min. Movements were recorded and analyzed using Any-maze software. The behavioral parameters included overall locomotion and time spent in different field zones. The percentage of time spent in each zone was calculated as (time in the zone/ total test duration) × 100.

### 2.9 Geomx DSP spatial transcriptomics

Formalin-fixed paraffin-embedded (FFPE) sagittal brain sections (5µm) were mounted on SuperFrost Plus slides (Thermo Fisher Scientific, cat.1255015) and processed following the GeoMX digital spatial profiling (DSP) platform (Bruker Nanostring Technologies, Seattle, WA, USA) RNA protocol guidelines [38]. Spatial transcriptomic profiling was performed using the Mouse Whole Transcriptome Atlas (WTA) assay, which measures the expression of approximately 21,000 genes in the tissue. The assay uses a cocktail of in situ hybridization (ISH) probes that bind to mRNA targets within fixed tissue sections. ISH probes are conjugated to light-sensitive, UV-photocleaved DNA-barcoded oligos, which provide highly multiplexed transcriptomic data. For histology-guided region of interest (ROI) selection, adjacent sections were stained with anti-HIV-1p24 (catalog. sc-65918) and anti-Aβ (6E10; BioLegend, cat:803001) antibodies to identify regions of viral infection and amyloid-β deposition. This guided the ROI selection and annotations. Fluorescently labeled antibodies were then used to identify the cell types of interest in the adjacent sections for DSP analysis. The middle section was stained with fluorescently labeled antibodies against IBA1-AF594 (Cell Signaling Technology, cat. 48934), GFAP-AF488 (Invitrogen, Cat. 53-9892-82), and NeuN-AF 647 (Abcam, cat. Ab190565) to visualize microglia, astrocytes, and neurons, respectively, along with SYTO13 dye for nuclear staining of the cells. The slides were scanned using a GeoMX DSP (Bruker NanoString Technologies). The ROIs were selected based on pathology and annotated as regions that expressed Aβ only, HIV-1 only, both Aβ and HIV-1 (Aβ+HIV-1), and no pathology. A total of 51 ROIs (17 ROIs per slide) were selected from three slides, two from the HIV-1-infected hNAIL mice group and one from the NOG group, with anatomically matched ROIs to one of the HIV-1-infected hNAIL mice. The UV-photocleavable DNA barcoded oligos from each segment were collected in individual wells of a 96-well plate. Once all tissue samples were processed and collected, next-generation sequencing (NGS)-based quantification was performed on them. The NGS libraries were sequenced using an Illumina NextSeq 550 instrument. Illumina FASTQ sequencing files were processed using the GeoMx NGS Pipeline version 2.3.3.10 to generate digital count conversion (DCC) files, which were subsequently analyzed using the GeoMx DSP Analysis Suite version 3.0 to perform data QC and Q3 normalization using all target groups. Data processing and quality control were performed using the GeoMX DSP analysis suite (v2.3.1) following the manufacturer’s recommendations. Differential gene expression was tested using an unpaired two-tailed t-test to compare different pathological regions (Aβ only, HIV-1 only, Aβ+HIV-1, and no pathology controls). Genes with log_2_ fold change> 0.5 and -log_10_ p-value ≥1.3 were considered differentially expressed genes (DEGs).

### 2.10 Statistical analyses

Statistical analyses were performed using GraphPad Prism v10.6.1. Group comparisons were assessed using one-way analysis of variance (ANOVA), followed by Tukey’s post hoc test. Spatial transcriptomic comparisons were performed using an unpaired two-tailed t-test in the GeoMx DSP Data Analysis suite. Statistical significance was set at p < 0.05.

## Results

### 3.1 hNAIL mice express the human APP mutation and support productive HIV-1 replication

To characterize hNAIL in HIV-1-AD pathologies, we evaluated human APP and Aβ expression, plasma viral load, and human cell reconstitution. NAIL mice, 2-3 days after birth, were transplanted with CD34⁺ HSC to generate human immunocytes. Successful engraftment was confirmed at 4 months by the presence of human CD45 + cells. Mice were then challenged with HIV-1_ADA_ for 8 weeks before being sacrificed (**Fig. 1A**).

**Figure 1:**
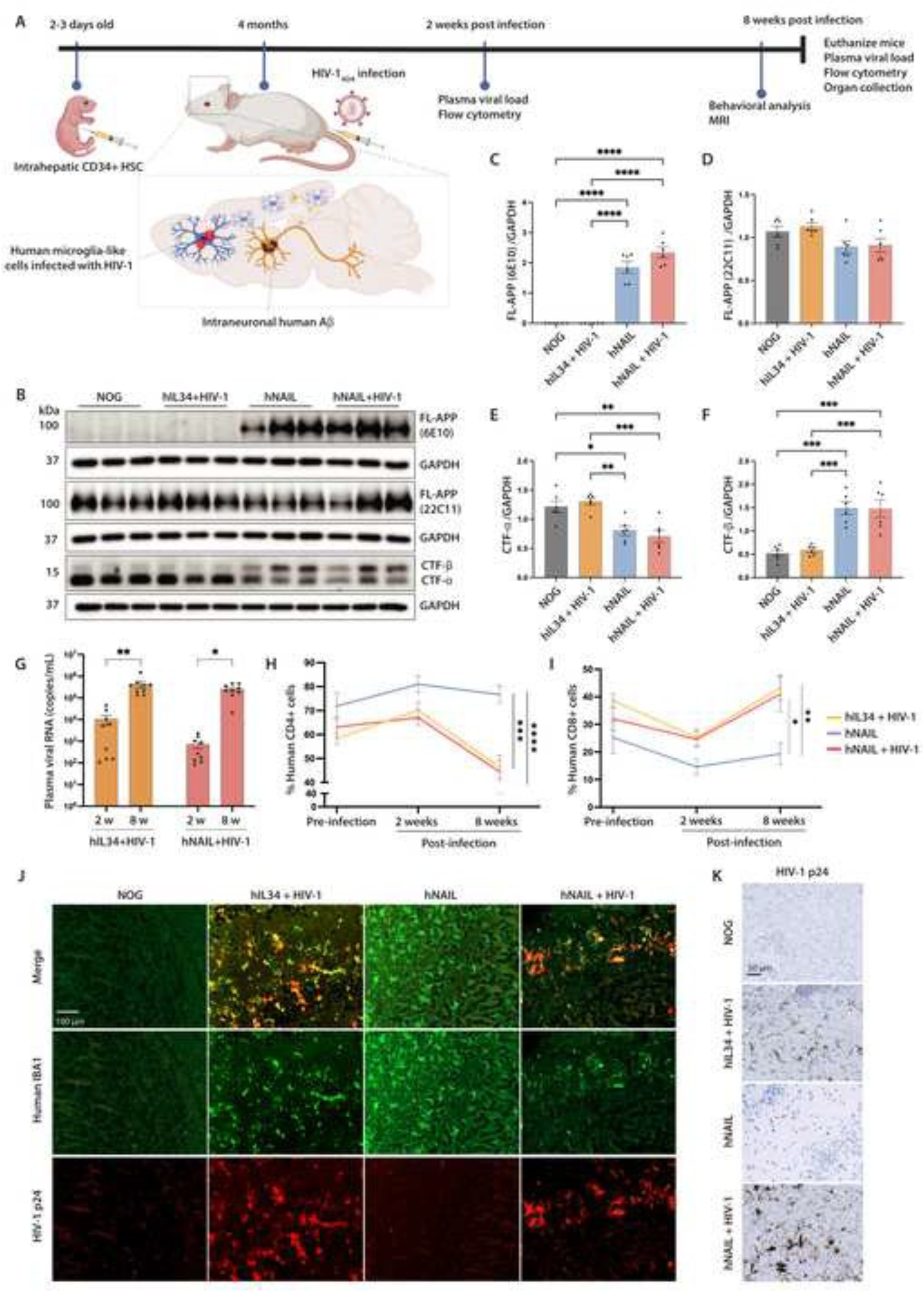
Characterization, humanization, and HIV-1 infection of the hNAIL mice. (A) Schematic overview of the experimental design and timeline for humanization, HIV-1 infection, behavioral analysis, MRI, and tissue collection. (B) Representative immunoblots of human-specific (6E10) or human/mouse-specific (22C11) full-length APP (FL-APP) and APP C-terminal fragments (CTF-α and CTF-β) from cortical lysates of 6-month-old NOG, HIV-1-infected hIL34, hNAIL, and HIV-1-infected hNAIL groups collected 8-weeks post infection, with GAPDH as the loading control. Brain samples were collected 8-weeks post-infection. Quantification of (B) human-specific FL-APP (6E10), (C) human/mouse non-specific FL-APP (22C11), (E) CTF-⍺, and (F) CTF-β fragments, normalized to GAPDH. (G) Plasma HIV-1 viral RNA levels pre-infection, 2 weeks, and 8 weeks post-infection. Flow cytometric analysis of (H) % human CD4+ cells and (I) % human CD8+ cells pre-infection, 2 weeks, and 4 weeks post-infection. (J) Representative IF of 5 μm thick sagittal brain sections showing co-localization (merge) of HIV-1 p24 (red) with human microglia-like cells (green) (hIBA1), indicating HIV-1 infection of human microglia-like cells. Scale bar = 100 µm (K) Representative IHC of 5 μm thick sagittal brain sections showing HIV-1 p24. Scale bar = 50µm. Data shown as mean ± SEM (n = 6-9 mice per group, both sexes). Statistical significance was determined using one-way ANOVA followed by Tukey’s post hoc test. *p < 0.05, **p < 0.01, ***p < 0.001, ****p < 0.0001.

Western blots were performed using a human-specific APP antibody (6E10), confirming the expression of mutant human APP in hNAIL and HIV-1-infected hNAIL mice (**Fig. 1B, C)**. No APP signal was detected in the NOG or HIV-1-infected hIL34 controls. Total APP was detected using 22C11 antibodies, which recognize both mouse and human APP, and was similar across all groups (**Fig. 1B,D**). Analysis of APP C-terminal fragments confirmed mutation-driven APP processing and increased CTF-β and reduced CTF-α levels in hNAIL and HIV-1-infected hNAIL mice (**Fig. 1B,E,F**). HIV-1 infection does not alter human APP expression or processing. This confirmed that CTF-β in hNAIL mice was driven by the knocked-in APP Swedish mutation, which enhances β-secretase cleavage of APP (**Fig. 1 B,C,F**). Next, we assessed HIV-1 replication in hNAIL mice. Plasma viral RNA levels increased between 2 and 8 weeks after infection in both HIV-1-infected hIL34 and HIV-1-infected hNAIL mice **(Fig. 1G)**. Flow cytometry analysis of peripheral blood revealed depletion of human CD4+ T cells and expansion of CD8+ T cells following infection **(Fig. 1H,I)**. Both were consistent with a productive HIV-1 infection.

We then evaluated HIV-1 infection in microglial cells in the hNAIL mice. Immunofluorescence staining (IF) performed on 5μm thick FFPE sagittal brain sections using antibodies against human-specific IBA1 (hIBA1) and HIV-1 p24 protein showed co-localization of HIV-1p24 with human microglia-like cells in both HIV-1-infected hIL34 and HIV-1-infected hNAIL mice **(Fig. 1J)**. HIV-1p24 immunohistochemistry (IHC) further confirmed HIV-1 infection in the HIV-1-infected hNAIL and HIV-1-infected hIL34 mouse brains (**Fig. 1K**). No viral protein was detected in uninfected hNAIL mice. No hIBA1 staining was observed in NOG control mice (**Fig. 1 J,K**). Successful engraftment of human microglia-like cells was further confirmed by IHC staining with human P2RY12 and TMEM119. Brain sections from HIV-1-infected hIL34, hNAIL, and HIV-1-infected hNAIL mice demonstrated immunoreactivity for human P2RY12 and TMEM119, indicating the presence of engrafted human microglia-like cells (**Supplementary Fig.1**). In contrast, NOG control mice lacked staining for both human P2RY12 and TMEM119.

Together, these results demonstrate that hNAIL mice express mutant human APP at physiological levels, exhibit increased CTF-β processing, and support disseminated (brain and systemic) HIV-1 replication. The hNAIL mice develop human microglia-like cells that are susceptible to HIV-1 infection, providing a novel platform for mechanistic studies of HIV-AD comorbidities in mice.

### 3.2 HIV-1 infection increases amyloid accumulation

To determine whether HIV-1 infection alters amyloid accumulation in the brain, soluble and insoluble Aβ_42_ levels were quantified in frozen hippocampal and cortical tissue samples. Brain tissue from NOG, HIV-1-infected hIL34, hNAIL, and HIV-1-infected hNAIL mice was collected at 8 weeks post-infection, and the amyloid load was quantified using Aβ_42_ ELISA. In the hippocampus (**Fig. 2A,B**), low levels of soluble human Aβ_42_ were detected in the control groups (NOG mice and HIV-1-infected hIL34), likely reflecting a defined background in the absence of the APP Swedish mutation. In contrast, hNAIL mice exhibited significantly higher soluble Aβ_42_ levels, driven by the presence of the APP mutation. Importantly, HIV-1 infection further increased soluble Aβ_42_ levels in the infected hNAIL mice compared to uninfected hNAIL animals, indicating that HIV-1 infection exacerbates hippocampal Aβ deposition. A similar pattern was observed in the cortex **(Fig. 2A,B),** where soluble Aβ_42_ levels were low in the control groups, increased significantly in hNAIL mice, and were further elevated by HIV-1 infection in hNAIL mice.

**Figure 2.**
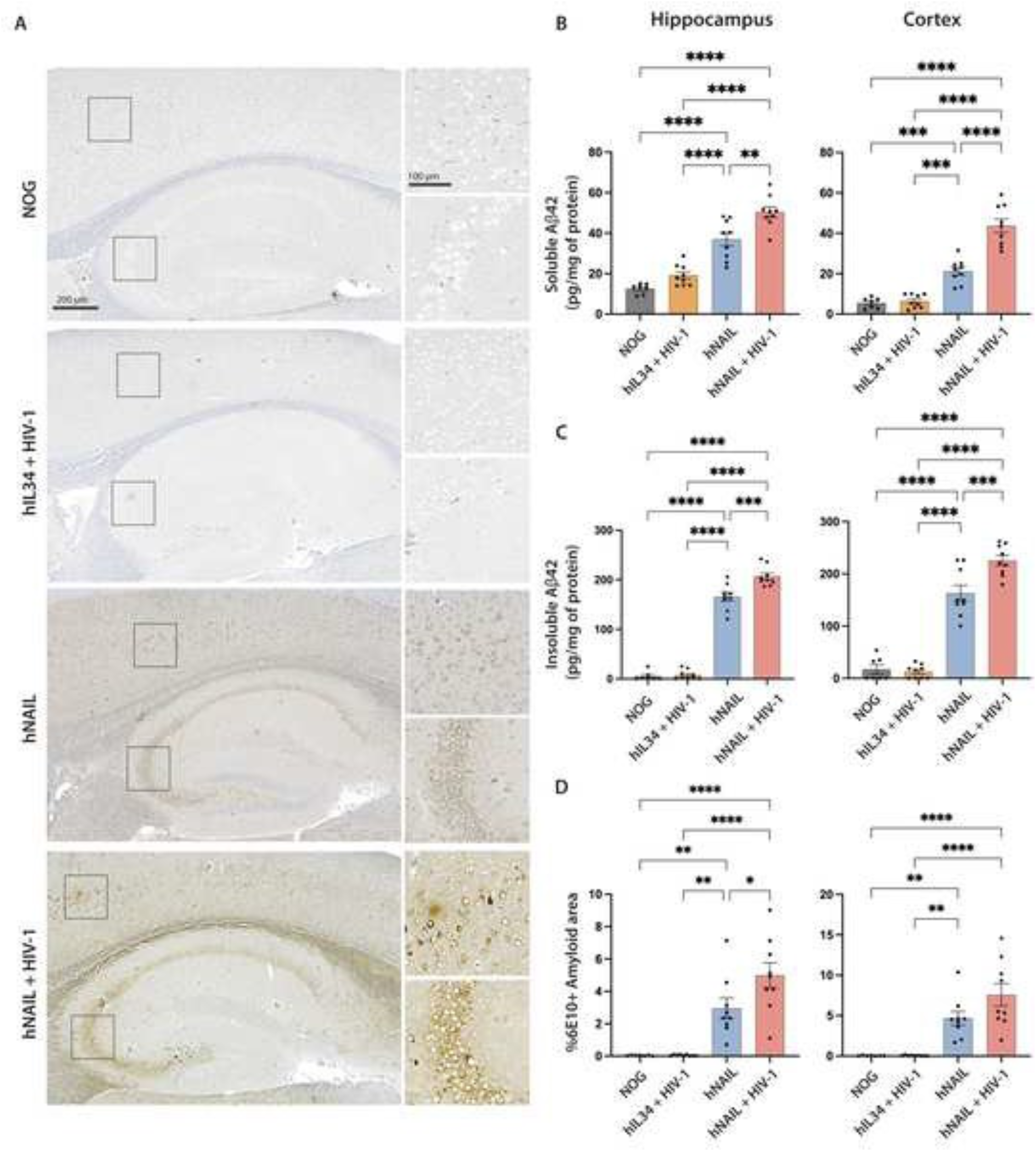
HIV-1 infection exacerbates amyloid pathology. (A) Representative IHC staining of 5 μm thick sagittal brain sections from 6-month-old NOG, hIL34+HIV-1, hNAIL, and hNAIL+HIV-1 mice collected 8 weeks post-infection, showing 6E10-positive human amyloid deposition in cortex and hippocampus. Insets highlight regions of amyloid accumulation. Scale bars: main image = 200 μm; insets =100 μm. ELISA quantification of (B) soluble human Aβ42 and (C) insoluble human Aβ42 from hippocampus and cortex lysates 8 weeks post infection. (D) Quantification of % 6E10-positive amyloid area in the hippocampus and cortex by IHC using automated image quantification, HALO software (Area Quantification v2.3.4 module). Data shown as mean ± SEM (n = 7-9 mice per group, both sexes). Statistical significance was determined using one-way ANOVA followed by Tukey’s post hoc test. *p < 0.05, **p < 0.01, ***p < 0.001, ****p < 0.0001.

Quantification of insoluble Aβ_42_ revealed a stronger APP-mutation-dependent phenotype. Insoluble Aβ_42_ levels remained low in the control groups (NOG and HIV-1-infected hIL34) in both the hippocampus and cortex **(Fig. 2A,C)**. In contrast, hNAIL mice exhibited marked accumulation of insoluble Aβ_42_ in both brain regions, which further increased following HIV-1 infection (**Fig. 2C**).

Quantitative IHC analysis using the 6E10 antibody, specific to human Aβ, was consistent with the ELISA findings. Automated whole-slide image analysis showed no detectable Aβ staining in the control groups (NOG and HIV-1-infected hIL34), whereas substantial amyloid accumulation was present in the cortex and hippocampus of hNAIL and HIV-1-infected hNAIL mice. In the hippocampus, HIV-1 infection significantly increased the 6E10-stained area in HIV-1-infected hNAIL mice compared to that in uninfected hNAIL controls **(Fig. 2D)**. In the cortex, there was a nonsignificant increase in the 6E10-stained area post HIV-1 infection compared to that in uninfected hNAIL mice. Notably, 6E10 immunostaining revealed predominantly intracellular amyloid accumulation in both the hippocampus and cortex, including in HIV-1-infected hNAIL mice. Together, Aβ_42_ ELISA and 6E10-IHC quantification demonstrated that HIV-1 infection increases both soluble and insoluble Aβ_42_ accumulation in the hippocampus and cortex, with a stronger effect on insoluble Aβ and increased intraneuronal Aβ accumulation.

### 3.3 HIV-1 infection increases amyloid-induced microglial activation

Neuroinflammation is a hallmark of both HIV-1 infection and AD pathobiology [39]. To assess whether increased amyloid deposition following HIV-1 infection was accompanied by glial activation, 5 μm sagittal brain sections were stained with the microglial marker IBA1 and the astrocytic marker GFAP. Quantitative analysis was performed using automated whole-slide image analysis software. In the hippocampus (**Fig. 3A,B**), activated microglia (defined as cells with less than 15 processes and a total process area of more than 15 µm^2^) were significantly increased in both HIV-1-infected groups (HIV-1-infected hIL34 and HIV-1-infected hNAIL) compared with NOG controls. Although the amyloid-only group (hNAIL) showed higher levels of reactive microglia than NOG mice, these differences were not statistically significant. Amyloid pathology alone resulted in approximately a twofold increase in reactive microglia in hNAIL mice, whereas the combined presence of Aβ and HIV-1 in HIV-1-infected hNAIL mice produced an approximately fourfold increase compared to NOG controls. These findings indicate that both amyloid pathology and HIV-1 infection contribute to microglial activation in the hippocampus (**Fig. 3B**). In the cortex **(Fig. 3C,D)**, differences in microgliosis were more pronounced. HIV-1-infected hNAIL exhibited the highest levels of activated microglia, with a significant increase compared to both NOG and hNAIL mice. The HIV-1 only group (HIV-1-infected hIL34) and amyloid-only (hNAIL) groups showed trends for increased reactive microglia when compared to NOG mice. However, the differences were not statistically significant **(Fig. 3D)**. These results indicate that HIV-1 infection enhances microglial activation in the hNAIL mouse brain, with a more pronounced effect observed in the cortex.

**Figure 3.**
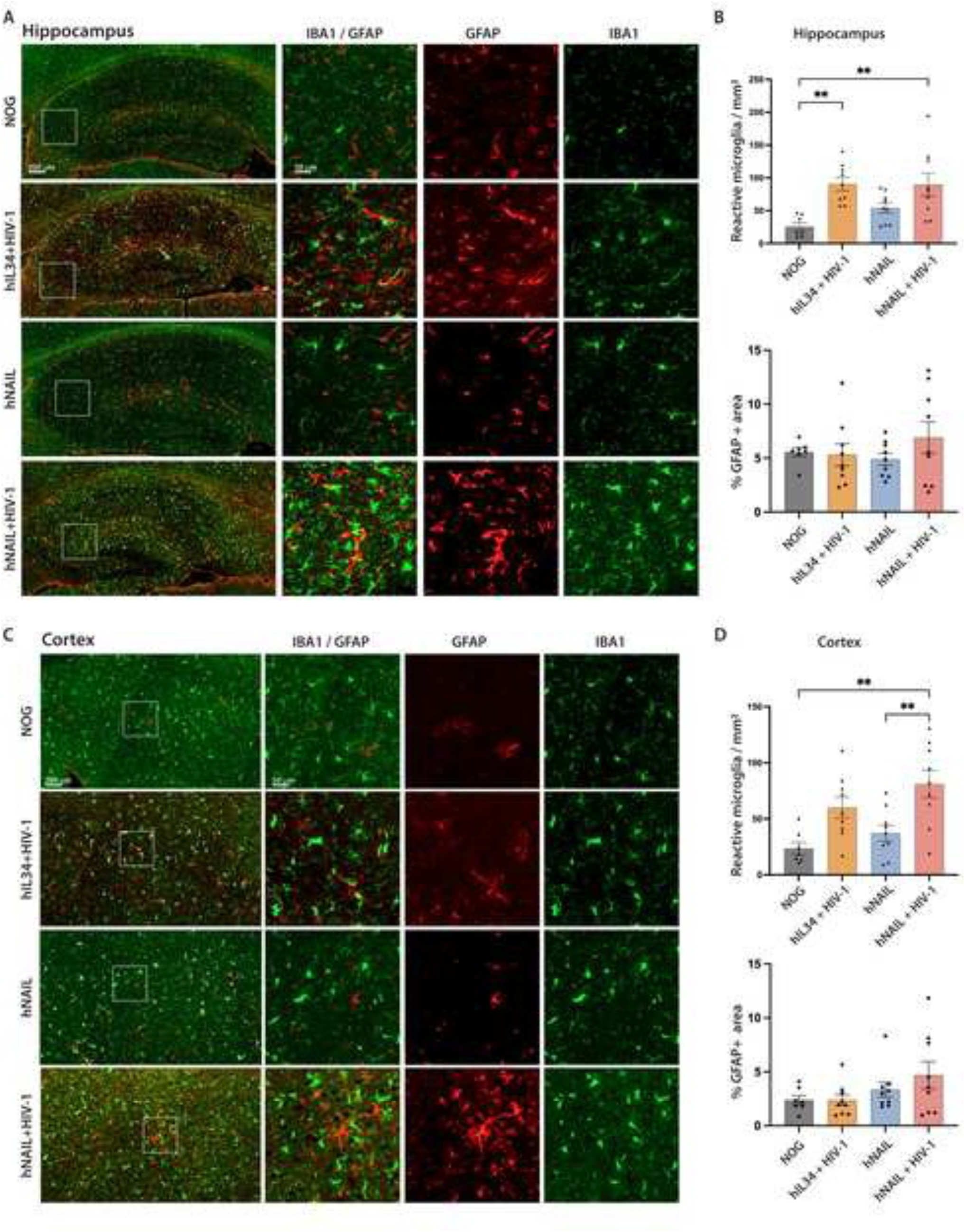
HIV-1 infection increases amyloid-induced microglial activation. Representative IF staining of 5 μm thick sagittal brain sections from 6-month-old NOG, hIL34+HIV-1, hNAIL, and hNAIL+HIV-1 mice collected 8 weeks post-infection, showing GFAP+ astrocytes (red) and IBA1+ Microglia (green) expression in the (A) hippocampus and (C) cortex. Insets show higher magnification with a single and merged channel. Scale bars: main image = 200 μm; insets =100 μm. Automated quantification of activated microglia in the (B) hippocampus and (D) cortex using HALO image analysis software, Microglia activation FL v1.0.6 module. Microglia with fewer than 15 processes and more than 15 μm^2^ cellular area were scored as activated. Automated quantification of % GFAP-positive area in the (B) hippocampus and (D) cortex using HALO image analysis software, Area Quantification FL v 2.3.4 module. Data shown as mean ± SEM (n = 7-9 mice per group, both sexes). Statistical analysis was performed using one-way ANOVA followed by Tukey’s post hoc test. *p < 0.05, **p < 0.01.

In contrast, astrocyte responses, measured as the percentage of GFAP-positive area, did not differ significantly among the groups in either the hippocampus or cortex (**Fig. 3A-D**). However, HIV-1-infected hNAIL mice exhibited a non-significant trend toward an increase in GFAP staining compared to other groups in both the hippocampus and cortex (**Fig. 3C,D**). The absence of widespread astrogliosis suggests that astrocytic responses may be more localized to the regions of infection rather than being broadly distributed across the tissue. This may also reflect differences in astrogliosis across different stages of HIV-1 infection and amyloid deposition. Collectively, these findings demonstrate pronounced microglial activation in the cortex and hippocampus of HIV-1-infected hNAIL mice, consistent with enhanced neuroinflammatory responses by combined HIV-1 infection and amyloid pathology. In contrast, astrocytic activation showed only a non-significant trend of increased % GFAP area, suggesting that astrocyte responses are localized.

### 3.4 HIV-1 infection exacerbates amyloid-induced loss of postsynaptic protein

Based on the increased amyloid accumulation and gliosis observed in HIV-1-infected hNAIL mice, we examined whether this was associated with alterations in synaptic proteins. To evaluate pre- and post-synaptic markers, 5μm sagittal brain sections were stained for synaptophysin and PSD-95 proteins for quantitative IF analysis. Synaptophysin, a presynaptic vesicle protein commonly used as a marker of presynaptic terminals, showed similar levels across all four groups in both the hippocampus and cortex (**Fig. 4A-D**), indicating no major changes in the presynaptic integrity. In contrast, PSD-95, a major scaffolding protein of the postsynaptic density that supports excitatory synapse stability and glutamatergic signaling, was significantly reduced in the groups expressing human Aβ. In the hippocampus, uninfected hNAIL mice exhibited a significant reduction in PSD-95 levels compared to NOG controls (**Fig. 4A, B**). The combined presence of amyloid pathology and HIV-1 infection (HIV-1-infected hNAIL) resulted in the lowest PSD-95 levels compared to the other groups.

**Figure 4.**
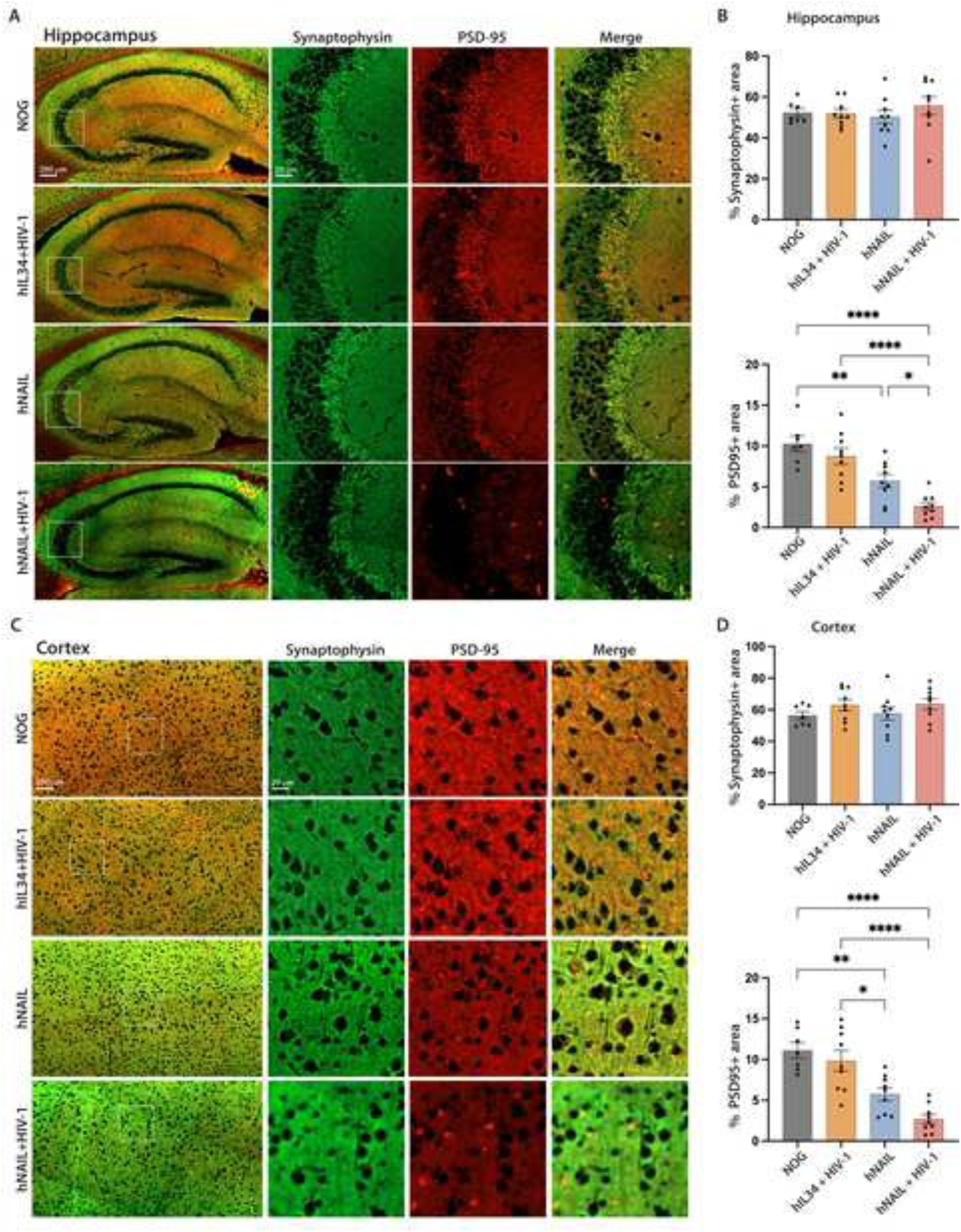
HIV-1 infection exacerbates synaptic loss in hNAIL mice. Representative IF staining of 5 μm thick sagittal brain sections from 6-month-old NOG, hIL34+HIV-1, hNAIL, and hNAIL+HIV-1 mice collected 8 weeks post-infection showing synaptophysin (green) and PSD-95 (red) expression in the (A) hippocampus and (C) cortex. Inset regions are shown in higher magnification with single and merged color channels. Scale bars: main image = 200 μm, insets = 20 μm. Automated quantification of % synaptophysin and % PSD-95-positive area in the (B) hippocampus and (D) cortex performed using HALO image analysis software, Area Quantification FL v 2.3.4 module. Data shown as mean ± SEM (n = 7-9 mice per group, both sexes). Statistical analysis was performed using one-way ANOVA followed by Tukey’s post hoc test. *p < 0.05, **p < 0.01, ****p < 0.0001.

A similar pattern was observed in the cortex, where PSD-95 levels were progressively reduced in hNAIL with or without HIV-1 infection compared to the NOG and HIV-1-infected hIL34 control groups (**Fig. 4C, D**). Although PSD-95 levels were lowest in HIV-1-infected hNAIL-infected mice, the differences following HIV-1 infection did not reach statistical significance in the cortex. These findings indicate that amyloid expression alone significantly reduces PSD-95 levels, whereas HIV-1 infection had a modest effect. The combined presence of amyloid pathology and HIV-1 infection resulted in the greatest reduction in postsynaptic density protein. Taken together, the selective reduction of PSD-95 and the absence of significant changes in synaptophysin suggest that postsynaptic structures are more vulnerable to amyloid-associated synaptic injury, with HIV-1 infection exacerbating postsynaptic loss. This selective loss of PSD-95 suggests an early disruption of the postsynaptic compartment rather than widespread degeneration of synaptic terminals.

### 3.5 HIV-1 infection amplifies amyloid-associated loss of neuronal integrity

To determine whether the increased amyloid accumulation, gliosis, and postsynaptic disruption observed in HIV-1-infected hNAIL mice were associated with neuronal injury, we evaluated neuronal integrity and cell loss using MAP2 and neuronal nuclei (NeuN) staining. Sagittal brain sections (5 µm) were stained for MAP2, a marker of neuronal structural integrity, and NeuN to assess neuronal density using quantitative IF analysis.

In the hippocampus (**Fig. 5A,B**), the MAP2-stained area was significantly reduced in HIV-1-infected hIL34, hNAIL, and HIV-1-infected hNAIL mice compared with NOG controls. The NOG mice exhibited long, continuous MAP2+ neuronal filaments, indicative of a preserved neuronal architecture **(Fig. 5A)**. In contrast, mice infected with HIV-1 only (HIV-1-infected hIL34), expressing human Aβ (hNAIL), or both pathologies (HIV-1-infected hNAIL) displayed truncated or fragmented MAP2+ neuronal processes, suggesting disrupted neuronal integrity. The reduced MAP2 staining area observed in the HIV-1-infected hIL34 and hNAIL groups indicates that either HIV-1 infection or amyloid pathology alone is sufficient to disrupt the neuronal architecture in the hippocampus. In the cortex, a similar reduction in MAP2 staining was observed in HIV-1-infected hIL34 and hNAIL mice compared to NOG controls, with the lowest levels detected in HIV-1-infected hNAIL mice **(Fig. 5C,D)**. However, unlike the hippocampus, the MAP2 staining area was significantly reduced in HIV-1-infected hNAIL mice compared to the hNAIL group, suggesting that HIV-1 infection contributes to an increased loss of neuronal structural integrity in the presence of amyloid pathology.

**Figure 5.**
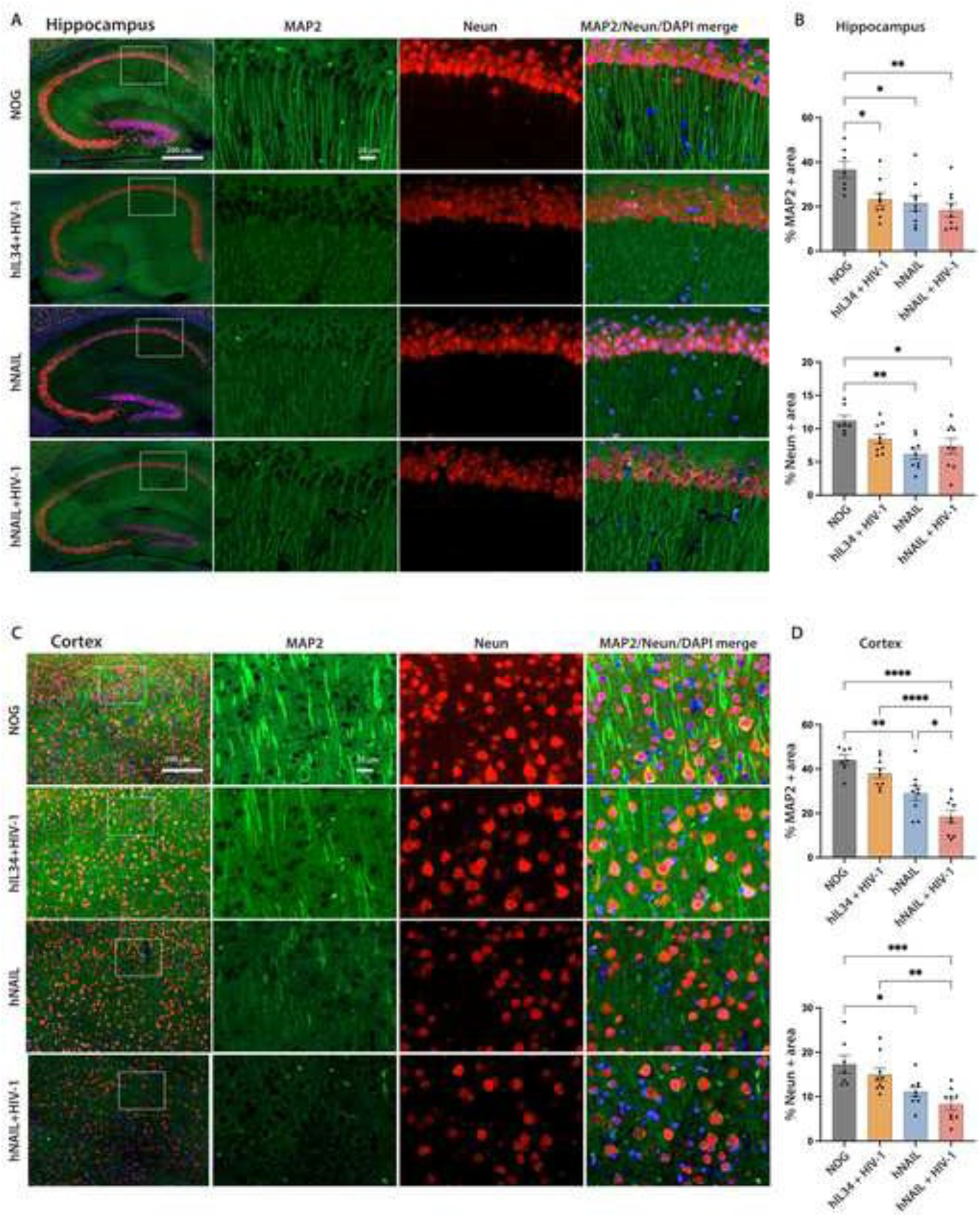
HIV-1 infection amplifies loss of amyloid-associated loss of neuronal integrity and density. Representative immunofluorescence staining of 5 μm-thick sagittal brain sections from 6-month-old NOG, hIL34+HIV-1, hNAIL, and hNAIL+HIV-1 mice collected 8 weeks post-infection showing MAP-2 (green) and Neun (red) expression in the (A) hippocampus and (C) cortex. Inset regions are shown in higher magnification with single and merged color channels. Scale bars: main image = 200 μm, insets = 20 μm. Automated quantification of % MAP2 and % Neun positive area in the (B) hippocampus and (D) cortex performed using HALO image analysis software, Area Quantification FL v 2.3.4 module. Data shown as mean ± SEM (n = 7-9 mice per group, both sexes). Statistical analysis was performed using one-way ANOVA followed by Tukey’s post hoc test. *p < 0.05, **p < 0.01, ***p < 0.001, ****p < 0.0001.

Further analysis of NeuN staining revealed a reduction in neuronal marker expression, primarily associated with amyloid pathology. In both the hippocampus and cortex, hNAIL mice with or without HIV-1 infection showed decreased NeuN-positive areas compared to NOG and infected hIL34 controls, indicating reduced neuronal density in the presence of amyloid pathology **(Fig. 5A-D)**. Notably, in the cortex, NeuN levels were significantly lower in HIV-1-infected hNAIL mice as compared to HIV-1-infected hIL34 mice, suggesting that the combined presence of HIV-1 and amyloid pathology contributes to higher neuronal loss (**Fig. 5C,D**).

Overall, these findings indicate that neuronal structural integrity can be disrupted by either HIV-1 infection or amyloid pathology alone in the hippocampus, whereas more significant neuronal injury is observed in the cortex when both factors are present. In contrast, reductions in NeuN-positive neurons were largely driven by Aβ and further exacerbated by HIV-1 infection, with a trend towards greater neuronal loss in the cortex following HIV-1 infection.

### 3.6 HIV-1 infection exacerbates amyloid-induced brain atrophy and behavioral deficits

To determine whether neuronal injury and loss observed in HIV-1-infected hNAIL mice were associated with structural brain changes and behavioral alterations, we assessed brain morphology using MRI and performed a battery of behavioral tests. Brain atrophy and structural changes were assessed using MRI 8-weeks post infection. Quantitative analysis revealed a significant reduction in the cortical area in HIV-1-infected hNAIL mice compared to NOG controls (**Fig. 6A, B),** indicating structural brain atrophy associated with combined amyloid pathology and HIV-1 infection. Cognitive performance was assessed using the novel-object recognition test. Both HIV-1-infected groups (HIV-1-infected hIL34 and HIV-1-infected hNAIL) showed a trend towards reduced preference for novel objects compared to NOG and hNAIL mice. However, these differences were not statistically significant (**Fig. 6C**). The hNAIL mice alone did not exhibit altered recognition memory. Working memory was evaluated using the Y-maze spontaneous alternation test, and no significant differences were observed among groups (**Fig. 6D**). Exploratory and anxiety-like behavior was assessed using the open field test. Notably, HIV-1-infected hNAIL mice spent significantly less time in the center compared to the NOG control mice, suggesting increased anxiety-like behavior in the combined pathology group (**Fig. 6E**). Other groups, HIV-1-infected hIL34 and hNAIL, also showed a reduced mean time spent in the center compared to the background NOG mice, although these differences did not reach statistical significance.

**Figure 6.**
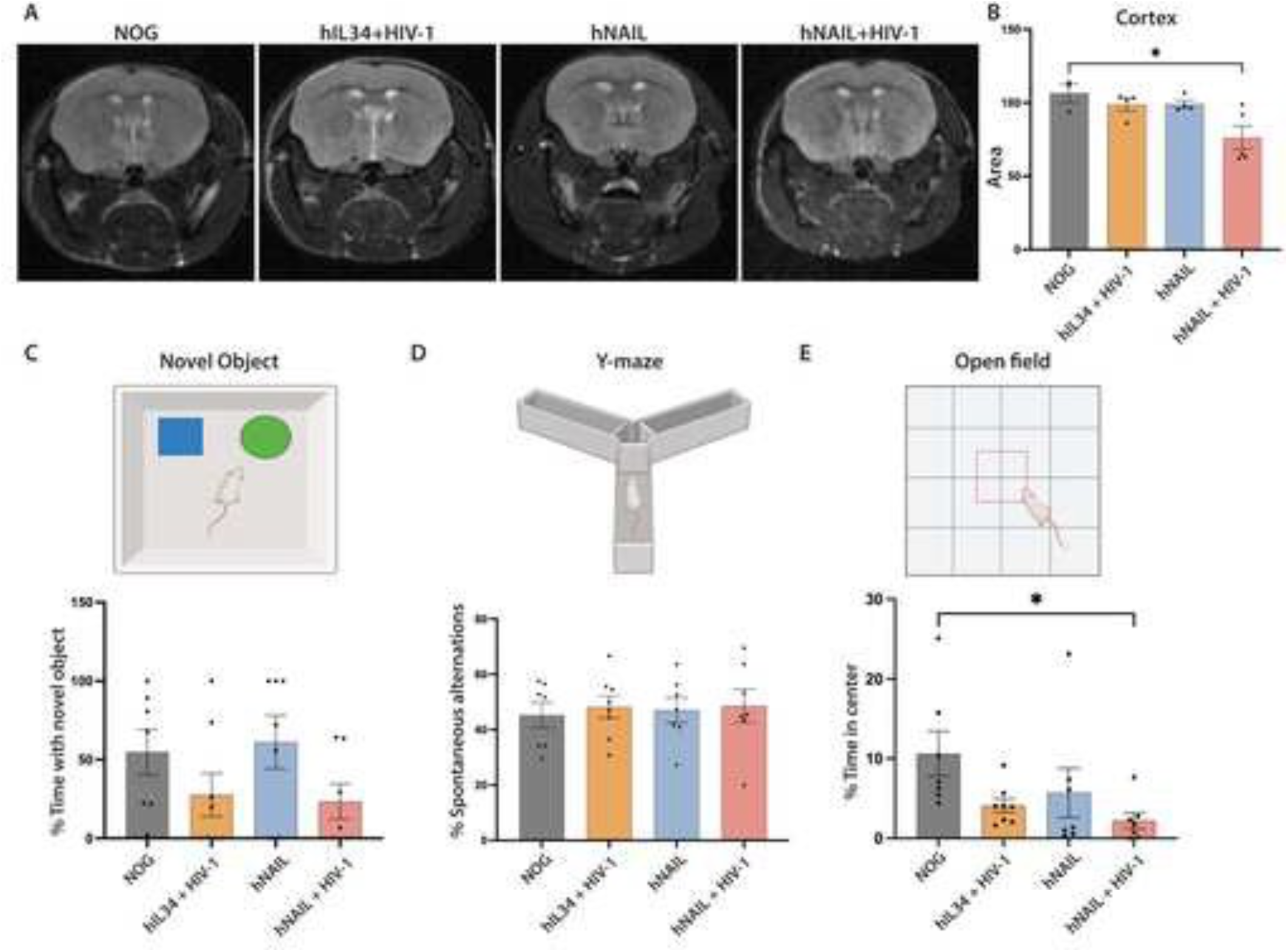
Brain structural and behavioral deficits following HIV-1 infection. (A) Representative coronal MRI scans of 6-month-old NOG, hIL34+HIV-1, hNAIL, and hNAIL+HIV-1 mice at 8 weeks post-infection (B) Quantification of the cortical area measured from MRI scans using ImageJ software. (C) quantification of Novel object test showing percentage time spent exploring the novel object. (D) Y-maze spontaneous alteration test measuring percentage spontaneous alternations as an indicator of spatial working memory. (E) Open field test showing percentage time spent in the center of the area as a measure of exploratory and anxiety like behavior. Data shown as mean ± SEM (n = 3-8 mice per group, both sexes). Statistical analysis was performed using one-way ANOVA followed by Tukey’s post hoc test. *p < 0.05.

Overall, these findings indicate that the combined presence of HIV-1 infection and amyloid pathology in HIV-1-infected hNAIL mice is associated with cortical atrophy and increased anxiety-like behaviors. In contrast, significant recognition and working memory deficits were not observed under the current experimental conditions.

### 3.7 Aβ and HIV-1 drive distinct transcriptional pathologies

Because amyloid plaques and HIV-1-infection occur in spatially restricted brain regions and involve multiple interacting cell types, we used spatial transcriptomics to resolve localized pathology-associated transcriptional changes that may be diluted in bulk transcriptomic analyses.

To define cell-type-specific transcriptional responses to Aβ, HIV-1, and their combination (Aβ+HIV-1), we performed GeoMx DSP spatial transcriptomic profiling on 5 μm thick FFPE sagittal brain sections. Regions of interest (ROIs) were classified and annotated as HIV-1 only, Aβ-only, Aβ+HIV-1, or no pathology control regions based on histopathological features observed in adjacent sections stained for HIV-1p24 or Aβ (6E10) **(Fig. 7A, supplementary 2A, 2B)**. The selected ROIs were processed using the NanoString GeoMx DSP platform (**Fig. 7A, supplementary 2C**). Differential expression was calculated relative to ROIs lacking both HIV-1 infection and amyloid pathology (no pathology control ROI).

**Figure 7.**
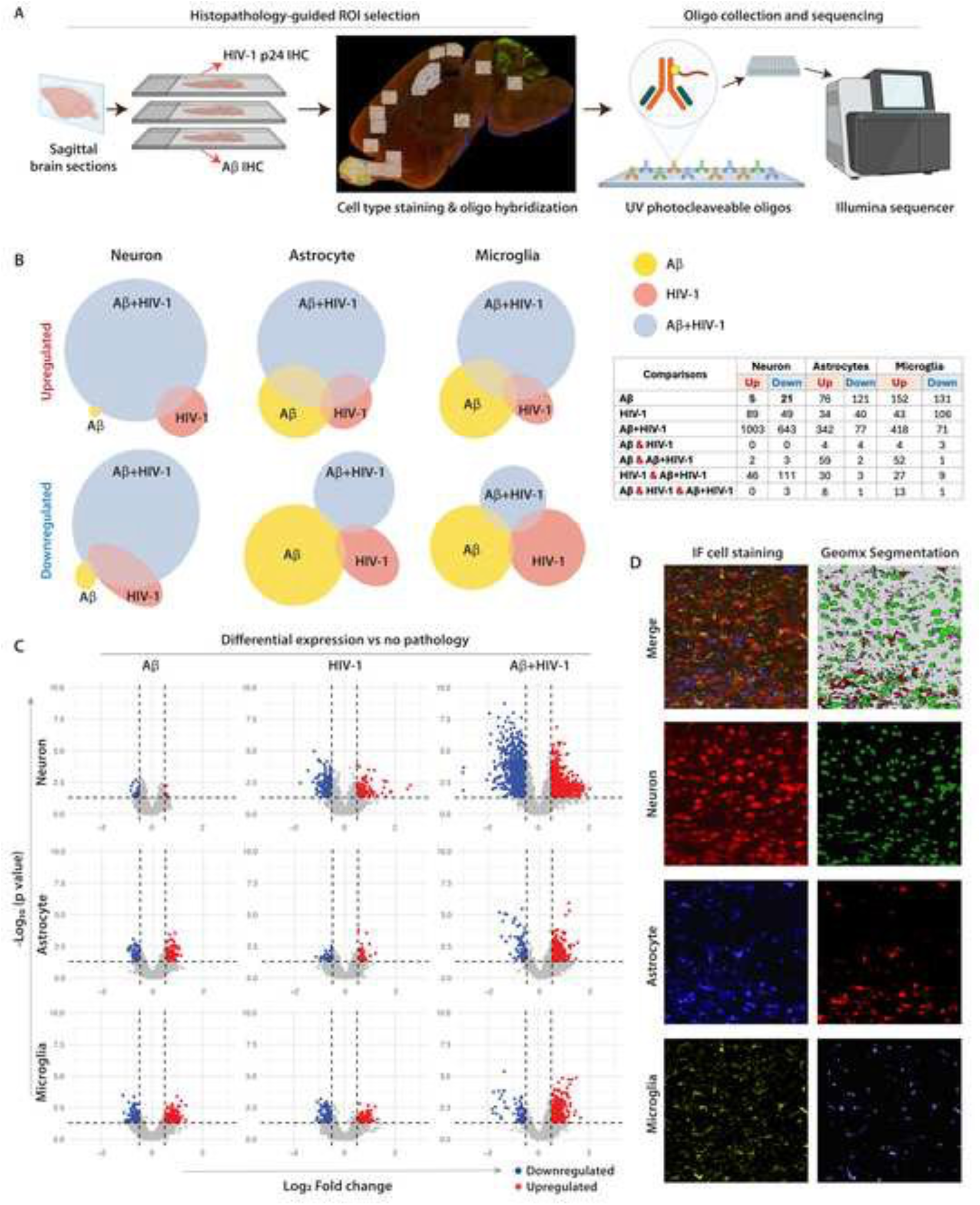
Aβ and HIV-1 drive distinct transcriptional changes, their combination induces strongest transcriptional changes. (A) Schematic overview of histopathology-guided ROI selection and NanoString GeoMx Digital Spatial Profiling (DSP) workflow. Adjacent sagittal brain sections were stained by IHC for HIV-1p24 and human Aβ (6E10) to guide ROI identification. The GeoMx DSP slide was immunofluorescently stained for cell-type markers for neurons (NeuN), microglia (IBA1), and astrocytes (GFAP) and hybridized with oligo-tagged RNA probes. The UV-photocleaved oligonucleotide tags from selected ROIs were collected for sequencing on an Illumina platform. (B) Venn diagrams showing the number of differentially expressed genes (DEGs) in neurons, astrocytes, and microglia across regions containing Aβ only, HIV-1 only, and combined Aβ+HIV-1 pathology compared with regions lacking both pathologies. The circle size is proportional to the number of DEGs. The accompanying table summarizes the number of upregulated and downregulated genes and their overlaps across the conditions. (C) Volcano plots showing DEGs for each cell type in Aβ only, HIV-1 only, and combined Aβ+HIV-1 regions compared to the control regions with no pathology. Each dot represents genes plotted by log_2_Fold-change (x-axis) and -log_10_p-value (y-axis). Red dots represent significantly upregulated genes, blue dots represent significantly downregulated genes, and gray dots indicate nonsignificant genes. (D) Representative IF staining and corresponding GeoMx spatial transcriptomic segmentation used for cell type identification, demonstrating concordance between IF-based cell labelling and GeoMx segmentation. The threshold for differential expression was set at log_2_FC ≥ 0.5 and -log_10_ (p-value) ≥ 1.3. Statistical analyses were performed using unpaired two-tailed t-tests.

Venn diagram analysis of DEGs from 34 ROIs across two brain sections from HIV-1-infected hNAIL mice, compared with no-pathology control ROIs, revealed largely distinct transcriptional signatures induced by Aβ and HIV-1 across all cell types (**Fig. 7B**). In neurons, no overlap (0 DEGs) was observed between Aβ- and HIV-1-regulated genes, indicating that each pathology drives an independent transcriptional program. In contrast, glial cells showed limited overlap in differentially expressed genes between Aβ and HIV-1 conditions, with microglia and astrocytes sharing seven and eight DEGs, respectively. Notably, regions with combined Aβ+HIV-1 pathology exhibited the largest number of differentially expressed genes across all three cell types. Neurons showed the strongest response (1646 DEGs), while astrocytes and microglia also demonstrated substantial transcriptional changes with 419 and 489 DEGs, respectively, indicating an amplified molecular response with dual pathologies. Consistent with these largely distinct transcriptional programs, the three-way overlap of Aβ, HIV-1, and Aβ+HIV-1 conditions was minimal, with only a small number of shared genes in glial genes and virtually no shared responses in neurons.

To further visualize the magnitude and distribution of transcriptional changes, we generated volcano plots comparing each pathological condition (Aβ, HIV-1, and Aβ+HIV-1) to the no-pathology control ROIs (**Fig. 7C**). Representative immunofluorescence labeling and GeoMx segmentation illustrated neuronal, astrocytic, and microglial compartments within each ROI to generate cell-type resolved differential gene expression (**Fig. 7D**). Immunofluorescence labeling for neurons (red), astrocytes (blue), and microglia (yellow) enabled cell-type identification within each ROI (**Fig. 7D)**. Aβ-only and HIV-1 only regions showed modest transcriptional changes, with relatively fewer genes above the significance thresholds (Log_2_FC ± 0.5 and -log_10_ p-value ≥ 1.3). In contrast, Aβ+HIV-1 regions demonstrated a higher number of significantly upregulated and downregulated genes in neurons, astrocytes, and microglia. This effect was particularly pronounced in neurons, where the combined pathology (Aβ+HIV-1) resulted in a greater number of differentially expressed genes compared to either Aβ or HIV-1 pathology alone (**Fig. 7C**).

Overall, these data demonstrate that HIV-1 infection and amyloid pathology drive largely distinct transcriptional programs, particularly in neurons, whereas glial cells exhibit limited overlap in their transcriptional responses. Notably, ROIs containing both Aβ and HIV-1 pathology showed the most extensive transcriptional alterations, indicating that dual pathology (Aβ+HIV-1) amplifies molecular responses beyond those induced by either condition alone.

### 3.8 Cell-type-specific neuroinflammatory signatures amplified in combined Aβ and HIV-1 brain subregions

To investigate the mechanistic basis underlying the amplified transcriptional changes observed in Aβ+HIV-1 brain regions, we performed a targeted analysis of DEGs identified by spatial transcriptomic profiling. Given the spatially restricted nature of the pathology and cell-type-resolved transcriptomic data, we focused on representative cell-type-specific genes rather than pathway enrichment analysis to better interpret localized molecular responses.

Before these comparisons were performed, an additional filtering strategy was implemented to minimize potential bias arising from the intrinsic transcriptional heterogeneity across brain regions, which is a known consideration in spatial transcriptomic analyses. To control for brain region-specific variability, ROIs were drawn from anatomically matched regions in a NOG background control mouse, mirroring the exact ROI annotations used in one of the HIV-1-infected hNAIL mouse slides. Differential gene expression analysis was performed between the corresponding ROIs in the NOG brain. Genes that were differentially expressed in both the HIV-1-infected hNAIL and anatomically matched NOG comparisons were removed. This filtering approach eliminated genes that could reflect intrinsic anatomical differences rather than pathology-driven, transcriptional changes. The remaining genes were considered pathology-specific and selected for targeted analysis.

In the targeted analysis of neuronal DEGs, genes associated with inflammatory signaling and innate immune signaling were significantly upregulated in Aβ+HIV-1 regions compared to the control ROIs, including IL-18, IFITM3, TRAF2, TRAF6, and TRIM26 (**Fig. 8A**). These genes are key components of NF-κB signaling and interferon-mediated innate immune pathways, indicating the activation of inflammatory transcriptional programs in neurons located within regions of dual pathology. Notably, IL18 has been shown to be a key modulator of CNS immune responses and has been associated with neuroinflammation and neurodegeneration [40]. These findings suggest that neurons exposed to combined Aβ and HIV-1 pathology adopt a heightened innate immune-associated inflammatory transcriptional profile.

**Figure 8.**
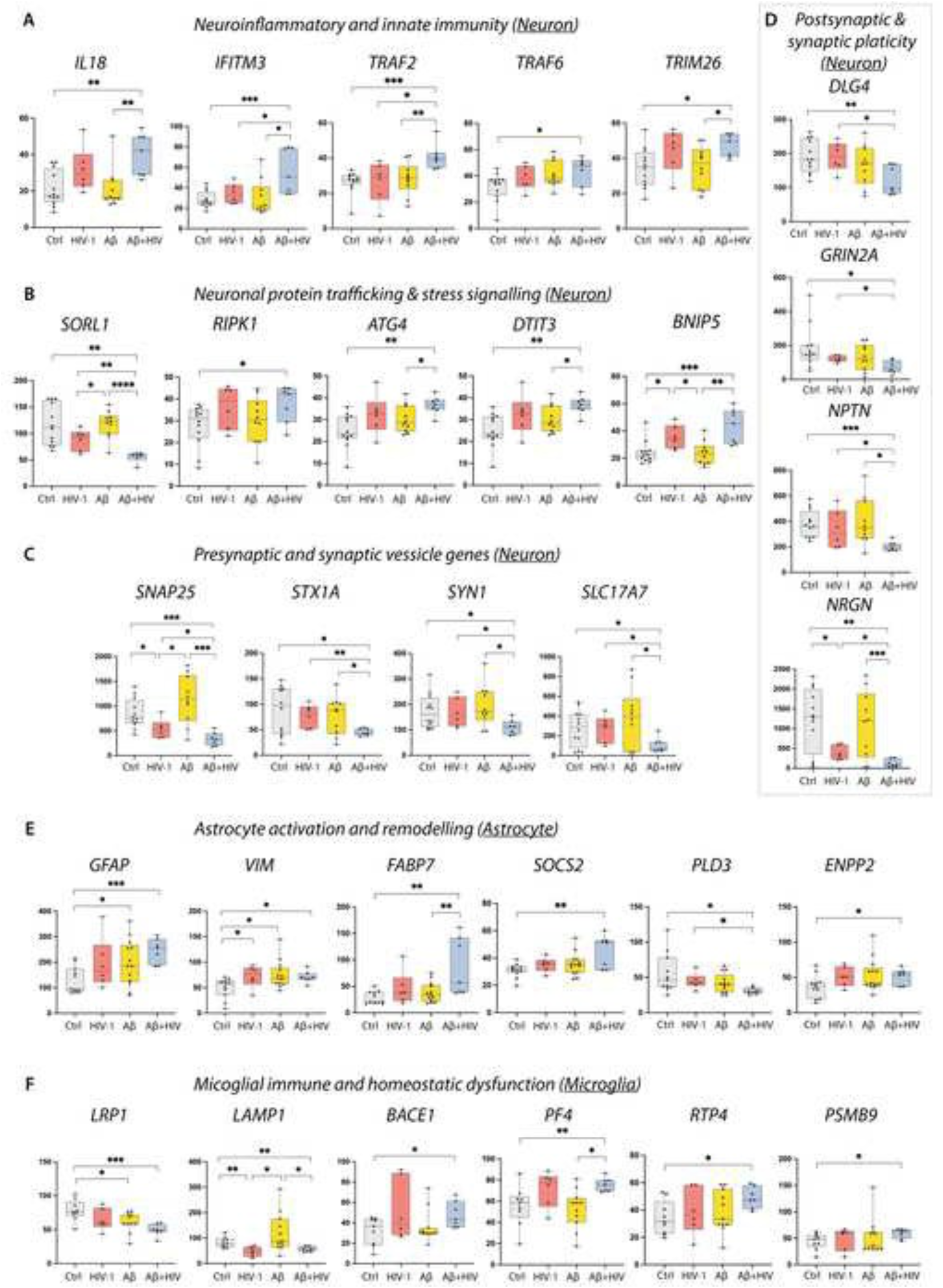
Aβ and HIV-1 drive distinct transcriptional changes amplified under dual pathology. Spatial transcriptomics was performed using Nanostring Geomx DSP on segmented neurons, astrocytes, and microglia from regions of interest (ROIs) containing no pathology (Ctrl), Aβ pathology alone (Aβ), HIV-1 pathology alone (HIV-1), and dual pathologies (Aβ+HIV-1). ROIs from two mice were merged for analysis, and DEGs overlapping with the control (NOG) mice were filtered before downstream comparison. (A) Neuronal inflammatory and innate immune genes (IL18, IFITM3, TRAF2, TRAF6, and TRIM26) are increased in Aβ+HIV-1 ROIs. B) Neuronal genes involved in protein trafficking and stress signaling (SORL1, RIPK1, ATG4, DDIT3, BNIP5) show altered expression in pathological conditions. (C) Presynaptic genes regulating vesicle docking and neurotransmitter release (SNAP25, STX1A, SYN1, and SLC17A7) are reduced in Aβ+HIV-1 regions. (D) Postsynaptic and synaptic plasticity genes (DLG4, GRIN2A, NPTN, and NRGN) were decreased in the Aβ+HIV-1 regions. (E) Astrocyte activation and remodeling genes (GFAP, VIM, FABP7, SOCS2, PLD3, and ENPP2) show altered expression across pathological conditions. (F) Microglial genes associated with immune regulation and homeostatic function (LRP1, LAMP1, BACE1, PF4, RTP4, and PSMB9) showed differential expression across conditions. Data are shown as box-and-whisker plots, with individual points representing each ROI. Boxes indicate the interquartile range with median; whiskers show the full data range. The Y-axis represents Q3 normalized gene expression. Statistical analysis was performed using an unpaired two-tailed t-test (*p < 0.05, **p < 0.01, ***p < 0.001, ****p < 0.0001).

Furthermore, neurons in dual pathology (Aβ+HIV-1) ROIs demonstrated altered expression of genes related to endosomal trafficking, cell organelle stress, and regulated cell death, including SORL1, RIPK1, ATG4, DTIT3, and BNIP5 (**Fig. 8B**). In particular, SORL1 regulates endosomal trafficking, including APP recycling, and its disruption is associated with endosomal changes and altered Aβ handling [41]. Concurrent increases in stress-related genes (DDIT3, ATG4) and cell death-associated signaling genes (RIPK1) indicate stress-linked pathology and autophagy activation in ROIs with dual pathology.

Given the neuronal loss and structural changes observed in Aβ+HIV-1 mice and that neurons displayed the highest number of DEGs in Aβ+HIV-1 regions, we further investigated the expression of neuronal synaptic genes to determine whether dual pathology is associated with alterations in the expression of synaptic proteins.

Analysis of neuronal synaptic genes revealed significant downregulation of multiple presynaptic components of the synaptic vesicle release machinery in Aβ+HIV-1 regions compared to no-pathology controls. Genes, including SNAP25, STX1A, SYN1, and SLC17A7, were significantly reduced in the dual pathology condition compared to single pathology (Aβ alone or HIV-1 alone) or no pathology ROIs (**Fig. 8C**). These genes encode proteins essential for synaptic vesicle docking, SNARE complex assembly, glutamate vesicle loading, and calcium-dependent neurotransmitter release. The significantly reduced expression of these presynaptic components suggests impaired vesicle fusion and diminished efficiency of glutamatergic neurotransmission in dual pathology ROIs compared to controls.

In addition to presynaptic alterations, key postsynaptic genes involved in excitatory synaptic organization and neuronal plasticity were reduced in the Aβ+HIV-1 regions. The expression of DLG4 (PSD95), GRIN2A, NPTN, and NRGN was significantly lower in dual pathology ROIs than in single or no-pathology controls (**Fig. 8D**). These genes play a central role in postsynaptic scaffolding density, NMDA receptor signaling, and stabilization of excitatory synapses. The downregulation of these postsynaptic genes indicates increased weakening of the synaptic architecture and reduced synaptic plasticity in neurons exposed to combined Aβ and HIV-1 pathology.

Astrocytes from the Aβ+HIV-1 regions also exhibited transcriptional changes consistent with reactive gliosis. Astrocytic activation and remodeling genes, including GFAP, VIM, FABP7, SOCS2, PLD3, and ENPP2, were differentially expressed in the Aβ+HIV-1 regions compared to the control region. The expression of GFAP and VIM, canonical markers of astrocytic activation and cytoskeletal remodeling, was increased in the Aβ+HIV1 region relative to the control regions (**Fig. 8E**). Similarly, FABP7 expression was higher than that in the controls, suggesting FABP7’s role in driving inflammatory responses in astrocytes in the region of dual pathology. Furthermore, SOCS2 expression was higher in the Aβ+HIV-1 regions than in the control regions. In contrast, PLD3 exhibited reduced expression, with the highest reduction observed in the dual Aβ+HIV-1 regions. ENPP2 showed a moderate increase in expression across pathological conditions, with higher levels in the Aβ+HIV-1 region. These changes indicate the presence of a localized reactive astroglial phenotype surrounding sites of dual pathology rather than global astrocytic activation across the brain.

Significant transcriptional changes were observed in genes associated with immune regulation and amyloid processing in microglia. These include LRP1, LAMP1, and BACE1 in the Aβ+HIV-1 regions (**Fig. 8F**). LRP1, a key receptor involved in Aβ clearance, showed the lowest expression in the dual-pathology ROIs. LAMP1 expression varied across conditions and was significantly reduced in Aβ+HIV-1 regions relative to Aβ alone, suggesting disruption of endo-lysosomal pathways. BACE1 expression was increased in the Aβ+HIV-1 regions, consistent with altered amyloid-related processing pathways under dual pathology. Additional genes, including PF4, RTP4, and PSMB9, also showed altered expression across pathological conditions, with the strongest shifts observed in the Aβ+HIV-1 regions. These genes are associated with immune signaling, synaptic regulation, and interferon-associated antigen processing, indicating changes in microglial immune activity and proteostatic pathways in regions with combined pathologies.

Together, this targeted analysis of representative cell-type-specific genes indicates that Aβ or HIV-1 alone produces relatively modest transcriptional changes, whereas dual pathology (Aβ+HIV-1) is associated with amplified and cell-type-specific transcriptional responses. Spatial transcriptomic analysis revealed increased expression of neuronal inflammatory genes, reduced expression of synaptic genes, astrocyte activation, and microglial immune dysregulation in Aβ+HIV-1 regions. These coordinated transcriptional changes across neurons, astrocytes, and microglia suggest that the co-occurrence of amyloid pathology and HIV-1 infection amplifies neuroinflammatory and neurodegenerative molecular genes in regions with dual pathology.

Notably, these localized transcriptional changes were not fully reflected in the global IHC quantification of presynaptic (synaptophysin) and astrocytic (GFAP) proteins. These findings suggest that localized transcriptional dysregulation of synaptic genes within Aβ+HIV-1 regions may precede or be diluted in global protein-level measurements, highlighting the complementary strengths of spatial transcriptomics and traditional histological approaches in resolving early molecular changes in neurodegeneration.

## Discussion

We used a novel humanized APP KI mouse model of early AD (hNAIL) that allows the development of human microglia-like cells. This model was effectively used to investigate the role of sustained HIV-1 replication in AD-associated neurodegeneration [34]. We showed HIV-1 infection exacerbates AD-like pathologies, including increased amyloid accumulation, microglial activation, synaptic and neuronal loss, brain atrophy, and behavioral alterations. Spatial transcriptomics revealed that Aβ and HIV-1 drive largely distinct transcriptional responses, which are amplified in the presence of dual pathology.

The hNAIL mice develop a sustained intracellular Aβ phenotype and do not develop aggressive extracellular plaque deposition [33, 42]. In contrast, AD models with rapid plaque formation often exhibit a high amyloid burden and neuroinflammation, which may obscure the effects of additional insults. Our findings demonstrate that HIV-1 infection increased Aβ accumulation in the brain, suggesting that viral infection may accelerate early AD-like pathology. These observations align with growing evidence that infectious agents and inflammatory stimuli can amplify amyloidogenic pathways [43, 44]. Although these studies examine diverse pathogens, they highlight a broader principle in which innate immune activation and host-pathogen interactions can modulate amyloid biology [45, 46]. Consistent with this notion, clinical and experimental studies have linked HIV-1 infection to altered amyloid metabolism. Autopsy studies have reported increased intraneuronal Aβ accumulation in HIV-1-infected individuals [11]. Clinical studies have shown increased Aβ levels in the brain and periphery, together with reduced cerebrospinal fluid (CSF) Aβ levels reflecting increased cerebral amyloid deposition [47].

In the present study, while HIV-1 infection alone led to increased microgliosis, the strongest activation was observed in HIV-1-infected hNAIL mice with combined HIV-1 and amyloid pathology. In HIV-1 infection, microglia and infiltrating macrophages serve as CNS viral reservoirs of HIV-1 and a driver of neuroinflammatory responses. Persistent microglial activation occurs even during suppressive ART [48]. In both AD and HIV-1-associated neurological disease, microglial responses can adopt multiple reactive states that evolve in response to limited pathologies [49]. Mechanistically, viral proteins have been shown to impair microglial phagocytosis and clearance of Aβ while promoting inflammatory signaling [50, 51]. Notably, the targeted spatial transcriptomic analysis of regions with combined Aβ and HIV-1 pathology revealed significantly increased expression of the amyloid-processing gene BACE1 and reduced expression of the amyloid clearance receptor LRP1. These findings suggest a shift toward enhanced amyloid deposition [52, 53]. In parallel, dysregulation of endo-lysosomal genes, such as LAMP1, further supports impaired microglial amyloid clearance [54]. Additional upregulated genes associated with interferon-mediated antiviral responses and immunoprotease activation, such as PF4(CXCL4), RTP4, and PSMB9, may further promote microglial inflammatory states that impair amyloid clearance and drive neuronal injury in regions with dual pathology [55-57].

Astrocytosis is a well-recognized component of HIV associated neurological disease. Astrocyte responses in our study exhibited a more spatially restricted pattern. While global IHC analysis revealed only modest and non-significant increases in the GFAP-positive area. Spatial transcriptomics identified localized upregulation of astrocyte reactivity and cytoskeletal remodeling markers such as GFAP and VIM in astrocytes within regions of combined pathology [58, 59]. Additionally, inflammatory signaling genes, such as FABP7, SOCS2, PLD3, and ENPP2, which have been implicated in AD pathology, were differentially expressed in regions of combined pathology, further supporting localized astrocytic activation [60-63]. Astrocytes often respond downstream of microglial activation in inflammatory CNS disorders, suggesting that the limited astrocytic response observed in this study may reflect an early stage of pathology [64]. Notably, only astrocyte and microglial DEGs showed partial overlap in Aβ+HIV-1 regions, whereas neuronal responses were largely distinct. This indicates shared glial transcriptional responses to distinct CNS insults, whereas neuronal responses to Aβ or HIV-1 appear largely nonoverlapping. This emphasizes investigating neuronal injury as a key downstream consequence of dual pathology.

Interestingly, spatial transcriptomic analysis revealed that neuronal transcriptional changes were most pronounced in regions with combined Aβ+HIV-1 pathology, whereas Aβ-only or HIV-1-only regions exhibited relatively limited neuronal DEGs. Histological analysis further demonstrated the highest loss of markers of neuronal integrity (NeuN), dendritic structure (MAP2), and post-synaptic density (PSD-95) only in animals with dual pathology, followed by the Aβ alone and HIV-1 alone groups. We previously demonstrated that intraneuronal amyloid accumulation drives neuronal loss in NA mice and that increasing the amyloid burden by introducing the PS1 AD mutation doubled neuronal loss [33]. While HIV-1 does not directly infect mouse neurons, the limited NeuN-positive neuronal loss observed is most likely due to inflammatory responses from infected microglia, macrophages, or lymphocytes. Nevertheless, HIV-1 replication in the brains of hNAIL mice facilitated by human microglia-like cells was sufficient to exacerbate intraneuronal amyloid-mediated neuronal injury, highlighting the synergistic impact of combined Aβ deposition and HIV-1 infection on neurodegeneration. Spatial transcriptomic analysis has revealed increased expression of neuroinflammatory and innate immune genes, such as IL-18, IFITM3 [65, 66], TRAF2, TRAF6, and TRIM26, in regions of dual pathology [67]. Notably, IL-18, a pro-inflammatory cytokine implicated in HIV-1-associated immune activation and AD-related neuroinflammation [40], was elevated with HIV-1 infection alone, but the highest levels were observed in dual pathology regions, suggesting it may represent a shared inflammatory pathway linking HIV-1-induced immune activation with AD-associated neurodegeneration.

Synaptodendritic injury is often observed in patients with HAND. MAP2 is linked to neuronal integrity and has been used as a neuronal injury marker in HIV-1-related neurocognitive impairments [68]. Synaptic dysfunction and loss are key neurological correlates of cognitive impairment in AD. Interestingly, IHC quantification revealed that the presynaptic marker, synaptophysin, remained relatively stable across groups. In contrast, the postsynaptic protein PSD-95 was reduced in hNAIL mice and further exacerbated by HIV-1 infection, whereas HIV-1 alone produced minimal changes. This suggests that postsynaptic compartments may be particularly vulnerable to the combined effects of Aβ and HIV-1-associated neuronal stress. PSD-95 (encoded by DLG4) plays a central role in organizing receptor complexes at excitatory postsynaptic synapses and maintaining synaptic plasticity and stability, and its reduction indicates impaired excitatory synapses [69].

Similarly, neuronal postsynaptic genes, such as GRIN2A, NPTN, and NRGN, which are involved in NMDA receptor signaling, neural adhesion, and synaptic plasticity, are reduced in regions with dual pathology [70, 71]. Dysregulation of NRGN expression has previously been linked to HIV-1-induced neuroinflammation and altered neuronal signaling, suggesting a mechanistic link between synaptic loss and cognitive dysfunction in HAND[72].

In spatial transcriptomic analysis, we detected downregulation of multiple presynaptic genes involved in vesicle docking and neurotransmitter release, including SNAP25, STX1A, SYN1, and SLC17A7, as well as a reduction in postsynaptic genes such as DLG4 [73, 74]. This suggests that synaptic dysfunction in dual pathology is both regional and cell type-specific, with earlier synaptic injury being spatially constrained before inducing widespread synapse loss.

Neurons in regions with combined Aβ and HIV-1 pathology exhibit coordinated disruptions in genes involved in protein trafficking and stress signaling. Notably, SORL1, a key neuronal sorting receptor that regulates APP trafficking and retromer-mediated endosomal recycling, is strongly reduced in Aβ+HIV-1 regions [41]. Concurrent increased expression of ER stress signaling (DDIT3), autophagy remodeling (ATG4), mitochondrial stress pathways (BNIP5), and inflammatory necroptotic signaling (RIPK1) suggests a coordinated neuronal stress response involving protein trafficking, organelle dysfunction, and inflammatory cell death pathways [75-77] Taken together, this suggests multi-cell organelle stress contributing to accelerated amyloid pathology and neuronal vulnerability in dual Aβ+HIV-1 regions.

Loss of brain volume is commonly used as a radiological imaging marker for AD [78]. We previously described neuronal loss-induced shrinking of cortical volume only in 12-month-old single APP-KI mice [33]. However, in this study, HIV-1-induced acceleration of neuronal loss resulted in a detectable cortical volume loss in 6-month-old HIV-1-infected hNAIL mice. Despite this, memory deficits were not significant, whereas reduced time spent in the center was significant, indicating anxiety-like behavior. This suggests early synaptic injury can produce detectable behavioral changes, even when memory deficits remain subtle.

The hNAIL model has several limitations. Behavioral changes were modest, likely due to limited cohort size and constraints of immunocompromised mice that restrict more sensitive assays [79]. The model primarily captures intraneuronal Aβ driven and does not progress to extracellular plaque, limiting assessment of late-stage plaque-associated neurodegeneration. To address this, next-generation KI models incorporating additional APP mutations are under development. Additionally, the relatively short infection timeline captures early interactions between HIV-1 and amyloid pathology, whereas both AD and HAND evolve over decades, necessitating longer-term studies to define chronic disease progression.

In summary, hNAIL mice provide a clinically relevant platform to study AD and HIV-1 comorbidity in a humanized immune context. The model captures early intraneuronal Aβ pathology and supports human microglia-like cells that can be infected with HIV-1, enabling the study of chronic infection alongside gradual neurodegeneration, similar to what is observed in PLWH [42]. It also offers a useful system to investigate how antiretroviral therapy (ART) influences amyloid processing and disease progression [80-82]. hNAIL model integrates knock-in AD pathology with a humanized, HIV-1 permissive immune system. This model provides a valuable platform for studying HIV-AD mechanisms and evaluating therapeutic strategies.

## List of abbreviations

AD: Alzheimer’s disease
Aβ: Amyloid beta
APP: Amyloid precursor protein
APP-KI: Amyloid precursor protein knock-in
ART: Antiretroviral therapy
ATG4: Autophagy related 4
BACE1: Beta-site APP cleaving enzyme 1
BNIP5: BCL2 interacting protein 5
CNS: Central nervous system
CTF: C-terminal fragment
DAB: 3,3′-Diaminobenzidine
DDIT3: DNA damage inducible transcript 3
DEG: Differentially expressed gene
DLG4: Discs large homolog 4 (PSD-95)
ELISA: Enzyme-linked immunosorbent assay
ENPP2: Ectonucleotide pyrophosphatase/phosphodiesterase 2
FACS: Fluorescence-activated cell sorting
FABP7: Fatty acid binding protein 7
FBS: Fetal bovine serum
FFPE: Formalin-fixed paraffin-embedded
GFAP: Glial fibrillary acidic protein
GeoMx: GeoMx digital spatial profiler
GRIN2A: Glutamate ionotropic receptor NMDA type subunit 2A
HAND: HIV-associated neurocognitive disorders
HIV-1: Human immunodeficiency virus type 1
HIV-1ADA: Macrophage-tropic HIV-1 ADA strain
HSC: Hematopoietic stem cell
IBA1: Ionized calcium-binding adapter molecule 1
IF: Immunofluorescence
IFITM3: Interferon-induced transmembrane protein 3
IHC: Immunohistochemistry
IL-34: Interleukin-34
IL18: Interleukin 18
KI: Knock-in
LAMP1: Lysosomal-associated membrane protein 1
LRP1: Low-density lipoprotein receptor-related protein 1
MAP2: Microtubule-associated protein 2
MAPT: Microtubule-associated protein tau
MRI: Magnetic resonance imaging
NAIL: NOG/ APP^KM670,671NL^/IL-34 mouse model
NeuN: Neuronal nuclei
NF-κB: Nuclear factor kappa-B
NMDA: N-methyl-D-aspartate
NOG: NOD.Cg-Prkdc^scid^Il2rg^tm1Sug^/JicTac
NPTN: Neuroplastin
NRGN: Neurogranin
P2RY12: Purinergic receptor P2Y12
PBL: Peripheral blood leukocyte
PF4: Platelet factor 4
PLD3: Phospholipase D family member 3
PLWH: People living with HIV
PSMB9: Proteasome 20S subunit beta 9
PSD-95: Postsynaptic density protein 95
PVDF: Polyvinylidene difluoride
RIPK1: Receptor interacting serine/threonine kinase 1
ROI: Region of interest
RTP4: Receptor transporter protein 4
RTN3: Reticulon 3
SLC17A7: Solute carrier family 17 member 7
SNAP25: Synaptosomal-associated protein 25
SNARE: Soluble N-ethylmaleimide-sensitive factor attachment protein receptor
SOCS2: Suppressor of cytokine signaling 2
SORL1: Sortilin-related receptor 1
STX1A: Syntaxin 1A
SYN1: Synapsin I
SYTO13: Nucleic acid stain SYTO13
TBST: Tris-buffered saline with Tween-20
TCID50: 50% tissue culture infectious dose
TMEM119: Transmembrane protein 119
TRAF2: TNF receptor-associated factor 2
TRAF6: TNF receptor-associated factor 6
TRIM26: Tripartite motif-containing protein 26
TSPO: 18-kDa translocator protein
VIM: Vimentin
WTA: Whole Transcriptome Atlas
hIL34: HSC humanized human IL-34 transgenic mice
hNAIL: HSC humanized NAIL mice
p-tau: Phosphorylated tau

## Acknowledgements

The authors would also like to thank the UNMC animal behavior core (RRID: SCR_018830) and the Bioimaging core (RRID: SCR_022481) for their assistance with the study. Illustrations were created using Biorender.com. We express our gratitude to the community for its continuous encouragement and support of this research. Lastly, we thank the Vice Chancellor’s Office of the University of Nebraska Medical Center for their core facility support.

## Consent statement

All human participants provided informed consent. All authors read and approve the final version of the manuscript.

## HIGHLIGHTS

- A first-in-kind humanized KI AD mouse model supports cell-tissue reservoir HIV-1 replication, which includes human microglia-like cells.
- HIV-1 brain replication increases Aβ levels and impairs synaptic and neuronal integrity.
- Spatial transcriptomic analysis shows that combined HIV-1 and Aβ pathology amplifies responses compared to either alone.
- Neurons are the most affected cell type in the brain by significant transcriptional changes linked to neuroinflammation and synaptic dysfunction.
- hNAIL mice provide a unique platform to study HIV-AD comorbidities, facilitating future therapeutic developments.

## RESEARCH IN CONTEXT

1. Systematic review: We reviewed prior clinical and experimental studies examining the relationship between HIV-1 infection and Alzheimer’s disease (AD)-like pathology. Although aged people living with HIV exhibit increased risk of cognitive impairment and AD-like features, mechanistic understanding remains limited due to the lack of models that support both productive HIV-1 infection and human-relevant amyloid pathology. Existing HIV models lack AD pathology, while current AD models do not permit sustained HIV-1 infection in a humanized central nervous system environment.
2. Interpretation: Our findings demonstrate that HIV-1 infection accelerates early AD-like pathology in a humanized knock-in model by increasing amyloid burden, enhancing microglial activation, and driving region- and cell type-specific transcriptional changes. Dual Aβ and HIV-1 pathology amplifies molecular injury signatures beyond either insult alone, with neurons emerging as particularly vulnerable despite not being directly infected.
3. Future directions: The hNAIL model provides a platform to define mechanisms linking chronic HIV-1 infection to AD-related neurodegeneration and to evaluate therapeutic strategies. Future studies will examine long-term infection, effects of antiretroviral therapy, and progressive amyloid pathology in next-generation models to better capture chronic disease progression.

## Supplementary files

Click here to access/download

Supplementary files

Supplementary materials .docx

